# Methods and techniques enabling multi-kilobase long-range genomic rewrite/replace editing

**DOI:** 10.1101/2023.08.05.551844

**Authors:** Christopher A Piggott, Laurentia V Tjang, Selina Chen, Utsav Tatu, Hannah Hirou, Antonina Jaroszewska, Emma McIntyre, Jocelin Chen, Joelle V Faybishenko, Richard V Gavan, Chris R Hackley

## Abstract

CRISPR enabled cell and gene therapies have the potential to revolutionize the field of genetic medicine. However, the vast majority of rare diseases remain untreatable due to the limitations of current tools and techniques. To date, most corrective therapeutic approaches have been restricted to mutation-by-mutation approaches, where either HDR, or newer techniques such as base or prime editing, rewrite small regions of DNA at a time (∼1-100 bp). While these approaches are powerful, short editing windows (relative to the size of human genes) are financially and/or technically incompatible with most rare-disease mutation profiles. Here, we demonstrate for the first time that CRISPR/Cas9 can be used to “rewrite” 7kb+ sections of the human genome simultaneously via a selection-free process we have named “long-range rewriting”. Long-range rewriting approaches are compatible with multiple nucleases, cell types and genomic loci, and can be used with both double-strand break (DSB) and non-DSB based approaches.

## Introduction

CRISPR and related techniques have proven immensely useful in various fields from basic R&D to Industrial and more recently therapeutic applications. The most basic use of CRISPR/Cas9 systems is to target a certain section of DNA in the genome and create small <u>In</u>sertions or <u>Del</u>etions (INDELs) to disrupt various genetic elements (protein coding sequences, promoters/enhancers, splice sites, etc).^1–4^ While a powerful technique, this approach is limited in its therapeutic applications due to the very limited diseases where “breaking” a gene results in a therapeutic benefit. Additionally, there has been a growing concern over unwanted local and global side-effects generated via deleterious double-strand break (DSB) mediated editing.^5,6^

As such, there has been a push to develop tools and techniques that enable more programmable and sophisticated editing while simultaneously reducing unwanted side-effects. These efforts have resulted in development of various Cas9 fusion proteins with expanded functionality such as Base and Prime editors^7,8^, CRISPRa and CRISPRi^9,10^, engineering of Cas proteins designed to manipulate endogenous repair pathway choice^11–13^, and a myriad of other functionalities^14^. These efforts have resulted in the ability to selectively replace specific bases, swap any base to another, tune gene expression without genomic modification, and manipulate repair pathways to enable more types of editing at high efficiency and reduced toxicity than previously possible. Additionally, new techniques and tools have been developed which allow for high efficiency insertions of large (10kb+) of DNA with reduced insertional side-effects^15,16^.

However, even with all this exceptional progress, a major gap still exists: the ability to perform multiple discrete simultaneous edits over thousands of base-pairs of DNA with high intervening regions of homology. There are many challenges to this approach. Unlike HDR with transgene insertions, simultaneous discrete mutations over kilobase windows requires extending HDR repair through regions of very high homology, which traditionally has been viewed as impossible, with editing being reduced to background levels ∼10-40bp away from the DNA cutsite^17^. In theory, traditional techniques such as HDR should allow for specific edits over this range, as endogenous homologous recombination proceeds over 100s of kilobases^18^. As such, we decided to investigate whether an HDR based approach using either Cas9-HR, a fusion protein designed to increase HR and decrease NHEJ pathway choice and wildtype Cas9 could be used to create targeted and simultaneous rewrites over thousands of base-pairs.

## Methods

### General PCR and cloning methods

DNA amplification was performed using PCR with PrimeSTAR GXL (Takara) using standard protocols and 35 cycles unless otherwise specified. Molecular cloning was performed using PCR to generate fragments with 15 bp overlapping ends and then assembling the fragments using In-Fusion Enzyme (Takara Bio) prior to transformation into chemically competent *E. coli*. Plasmid DNA for transfection was isolated from *E. coli* using the ZymoPURE Plasmid Miniprep Kit (Zymo Research Corporation) or PureLink Fast Low-Endotoxin Midi Plasmid Purification Kit (Thermo Fisher Scientific), both which remove bacterial endotoxins. Primers and sequences of constructed repair templates are shown in **Supplementary Table 1**.

### Cell culture conditions

H1299 and HEK293T cells were grown in RPMI-1640 (Corning) and Dulbecco’s Modified Eagle Medium (DMEM, Corning) each supplemented with 10% fetal bovine serum. Cells were maintained at 37°C with 5% CO_2_.

### Tissue culture transfection protocol and genomic DNA preparation

H1299 or HEK293T cells were seeded on 24-well plates. Two days post-seeding, cells were transfected at approximately 80% confluency with Lipofectamine 3000 Transfection reagent (Thermo Fisher Scientific) according to the manufacturer’s protocols and 500 ng of Cas9-HR or Cas9 plasmid, 300 ng of sgRNA plasmid, and 30-50 ng of PCR product repair template. Cells were cultured for 3-5 days post-transfection after which genomic DNA was extracted using the Quick-DNA Microprep Kit (Zymo Research Corporation).

### Next-generation DNA sequencing of genomic DNA regions of interest

Genomic regions of interest were amplified from genomic DNA by PCR. Primers and PCR conditions are listed in **Supplementary Table 1**. PCR products were then run on an agarose gel and the desired band was excised and purified using the GeneJET Gel Extraction Kit (Thermo Fisher Scientific). Purified PCR products were prepared for sequencing using the standard Illumina DNA Prep protocol and reagents (Illumina DNA Prep (M), IPB), and subsequently sequenced using the MiSeq (Illumina) and MiSeq Reagent Kit v3 (Illumina, MS-102-3001) using standard sequencing conditions.

Illumina BaseSpace was used for fastq generation from raw BCL files. Subsequently, fastqs were trimmed using Trimmomatic with the parameters of SLIDINGWINDOW:10:30 and MINLEN:60 for quality trimming and ILLUMINACLIP:NexteraPE-PE.fa:2:30:10:2:True for adapter trimming. The trimmed sequence data were then aligned using bowtie2 with their respective reference sequences extracted from GRCh38 via NCBI. Custom python code using the pysam library (samtools/htslib) was used to generate a pileup of the number of nucleotides read at each position in the reference sequence, as well as parse individual reads into a CSV format, including INDEL statistics per read.

Pileup data were then used to quantify editing rate by calculating the percentage of read nucleotides at a given position that matched the intended edit at that position. Two types of editing rate datasets were produced: background-subtracted and non-background-subtracted. Background subtracted datasets had their editing rates subtracted by the editing rate percentage from the respective control containing nuclease and repair template only for each replicate. Non-background-subtracted datasets used the raw editing rate calculated from the pileup data. Read-level data were filtered down to a 50 bp window at the cut site for each guide, +-25 bp flanking each side. A percentage was then calculated for reads overlapping this window, creating datasets with the number of INDELs at each cleavage area. These datasets were then averaged across replicates, again producing mean and standard error of the mean values.

## Results

### Two Guide HDR based strategies using both Cas9-HR and Cas9 can be used to introduce 12 simultaneous single base changes in ∼3.3kb editing window

A major limitation of HDR based multiplexed gene-writing is the very small editing window limited to 10-40 bp around the guide-cut site. To determine whether it is possible to increase the size of this window, we theorized that employing a two-guide strategy, where two guides cutting determines the editing window, combined with a linear dsDNA template to mimic natural homologous recombination templates could be used to “force” the endogenous recombination machinery to read all the way through the intended editing window, allowing for both dramatically extended window size and even edit distribution.

Our first target was exon 14 of the Factor VIII gene, a therapeutically relevant locus with over 65 individual mutations classified as severe or likely-severe for Hemophilia A in NCBIs ClinVar^19^. Initially H1299 (CRL-5803) cells were transfected with plasmids encoding Cas9-HR or Cas9, dual guides under the pU6 promoter, and a PCR generated linear dsDNA template. The repair template was designed to introduce 9 diagnostic mutations in the exon 14 coding sequence, roughly scattered throughout the ∼3.3kb editing window, with 4 mutations destroying intronic PAM sites at our identified guide sites (Fig 1A). After transfection, various time-points (1, 3, 5, 10 days) post-transfection were assayed via amplification and restriction digest, with day 3 showing the most consistent editing (data not shown). As days 2-3 are very common time points for other CRISPR editing assays, we chose to use day 3 for all subsequent experiments unless noted. Finally, though restriction digests offer a facile and inexpensive assay, we wanted to ensure our data was of the highest quality and informationally dense, and so moved to the NGS based protocol shown in Fig 1B. Initially, we designed 4 guide pairs, with two pairs (G2+G5, G2+G6) showing good editing activity across the window. We used standard bioinformatic tools to align quantify desired editing at intended locations. Surprisingly, we saw remarkably consistent editing rates across the entire editing window with both Cas9 and Cas9-HR (Fig 1C). Another guide pair also showed good editing levels, though Cas9-HR showed lower background and more consistent editing than Cas9 (Fig 1D). Analysis of NGS sequencing showed consistent INDEL generation for both G2 and G5, though G6 showed less (Fig 1E), with editing rates and INDEL results from all guide pairs shown in Figure 1S A and B. Again, we see INDEL dependent editing, though interestingly only one guide out of the pair seemingly needs to effectively produce INDELs for chunk editing to succeed, like seen in Fig 1D.

**Figure 1:**
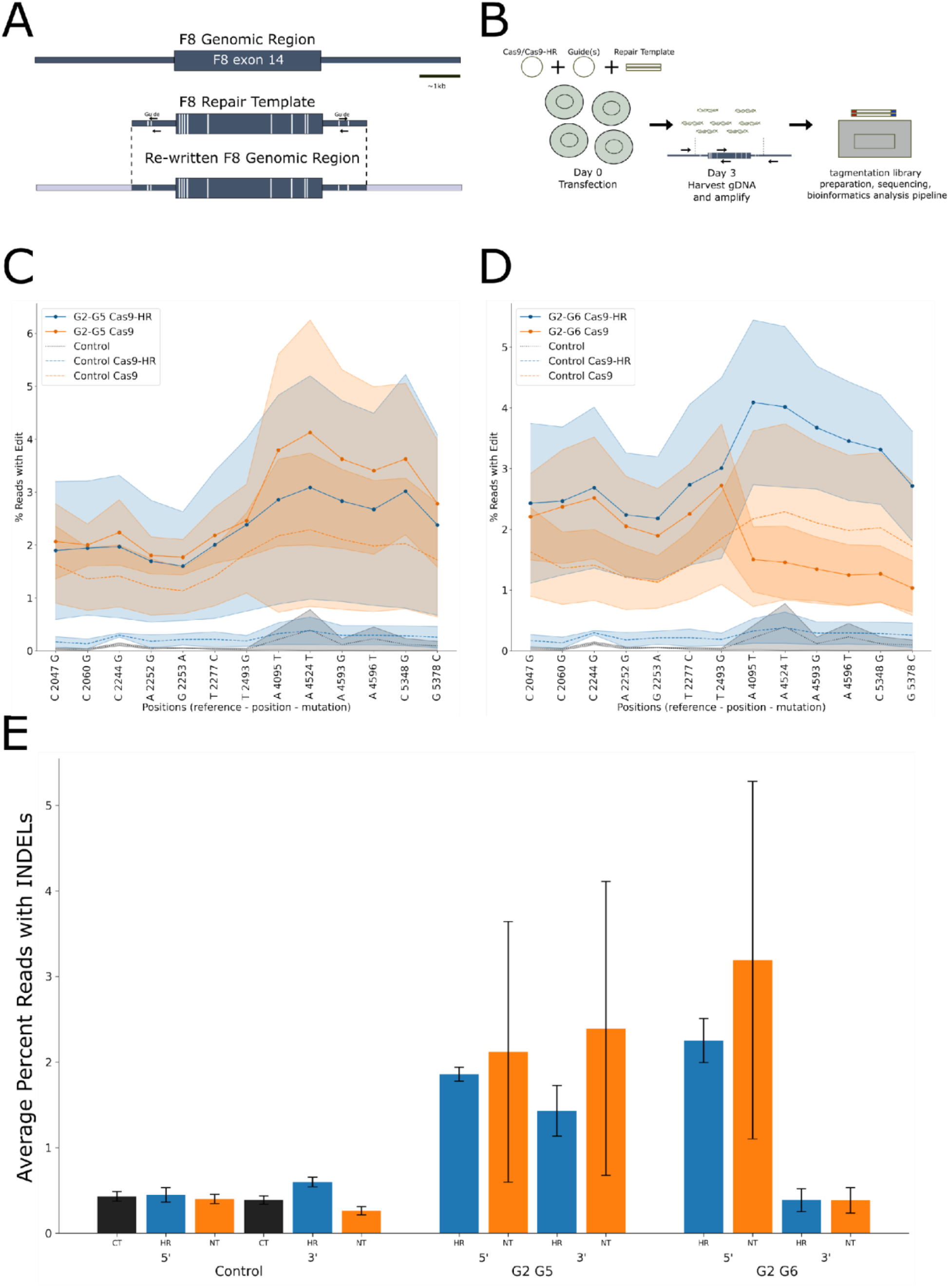
Cas9-HR and Cas9 be used to rewrite a ∼3.3 kb section of F8 exon 14. **1A**: Diagram depicting repair template design used to test rewriting capabilities. White bars denote intended single base changes, arrows indicate orientation of guides and approximate location in repair template/genome, dashed lines show end of repair template relative to genomic DNA. **1B:** Flow chart showing general editing strategy and downstream assays used for this and all subsequent figures. **1C**: Graph showing editing rates across 3+kb of genomic DNA using guide pair G2+G5. Cas9-HR, guides, and RT in blue; Cas9, guides, and RT in orange; Cas9-HR and RT dashed blue; Cas9 and RT dashed orange; untransfected control in dashed, black. Shading shows the SEM for each treatment, n=2-3 per treatment. **1D**: Graph showing editing rates across 3+kb of genomic DNA using guide pair G2+G6. Cas9-HR, guides, and RT in blue; Cas9, guides, and RT in orange; Cas9-HR and RT dashed blue; Cas9 and RT dashed orange; untransfected control in dashed, black. Shading shows the SEM for each treatment, n=2-3 per treatment. **1E**: Graph showing the average percentage of reads containing INDELs around +/- 25 of 5’ and 3’ cut sites. Error bars show SEM, n=2-3.

### Two guide editing strategies are necessary and sufficient to drive long-range editing in a second 4.5kb region

Given these results overturn over many years of precedent, we wanted to understand if the two-guide strategy was both necessary and sufficient to allow for long-range editing. Given our initial assay for F8 had non-trivial background, particularly in Cas9 samples, we decided to assay another locus, this time the region encompassing exons 6-8 in the PAH gene. We designed our repair template to contain three diagnostic mutations, one in each exon, and four PAM destroying mutations in the surrounding intronic regions (Fig 2A). Four different dual guide combinations were tested, with the best performing shown in Fig 2B. Again, both Cas9-HR and Cas9 show relatively consistent editing levels throughout the editing window, though this time Cas9 showed higher levels of editing than Cas9-HR, with background being very low for all types of controls.

**Figure 2:**
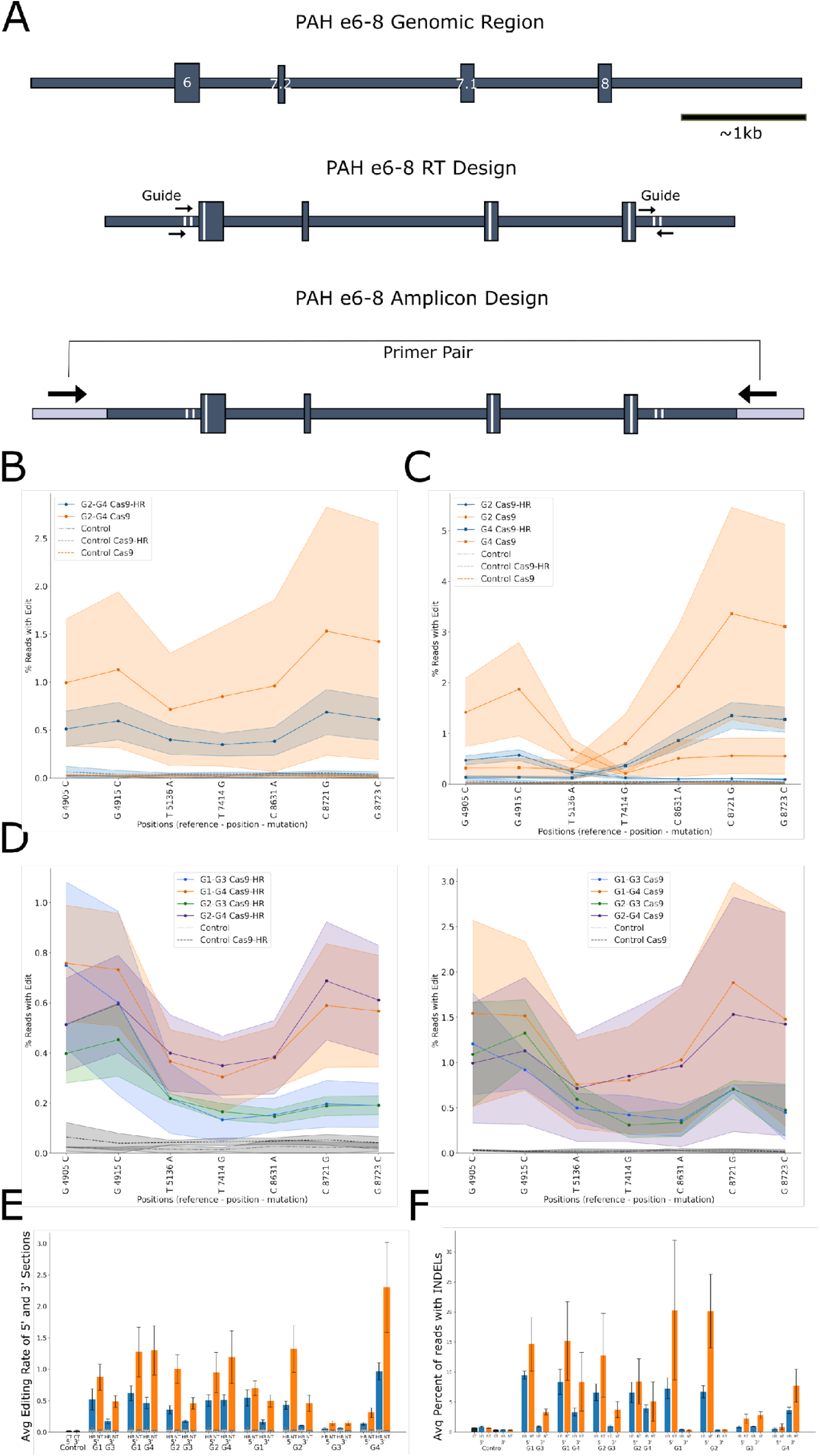
Two Guide strategies are necessary and sufficient to drive HDR based rewriting of ∼4.5kb region of PAH exons 6-8. **2A**: Diagram showing the structure of the ApoB exons 30 genomic region. The regions denoting the repair template are shown by dark blue shading, vertical white bars denote specific mutations, short black arrows (guides used and orientation), bottom shows integrated repair template with primers used for amplification from genomic DNA (large black arrows and grey shaded region respectively). **2B**: Graph showing editing rates across ∼4kb of genomic DNA using guide pair G2+G4. Cas9-HR, guides, and RT in blue; Cas9, guides, and RT in orange; Cas9-HR and RT dashed blue; Cas9 and RT dashed orange; untransfected control in dashed, black. Shading shows the SEM for each treatment, n=2-3 per treatment. **2C**: Graph showing editing rates across ∼4kb of genomic DNA using either single guides G2 or G4. Cas9-HR, guides, and RT in blue; Cas9, guides, and RT in orange; Cas9-HR and RT dashed blue; Cas9 and RT dashed orange; untransfected control in dashed, black. Shading shows the SEM for each treatment, n=2-3 per treatment. **2D**: Graph showing guide dependent effects of editing for four distinct primer pairs: G1+G3, G1+G4, G2+G3, G2+G4. Left, Cas9-HR; right, Cas9. **2E**: Average editing of various double or single primer pairs divided into 5’ and 3’ regions. Blue, Cas9-HR; Orange, Cas9; Black, untransfected control; Grey hatched bars, Cas9-HR or Cas9 and no guide RT controls. **2F**: Graph showing the Average percentage of reads containing INDELs around +/- 25 of 5’ and 3’ cut sites for single and double guide combinations. Error bars show SEM, n=2-3.

We then wanted to test whether two guide combinations were necessary to achieve long-range editing, so we transfected cells with only one guide and the same repair template (Fig 2C). Unsurprisingly, we saw a dramatic drop in editing efficiency outside of the traditional ∼10bp window (though levels were still generally above background at ∼100bp). Interestingly, the single guides showed both higher editing and INDEL efficiency at both 5’ and 3’ ends, likely indicating in our experiments that levels of nuclease may likely be limiting when multiple guides are added. Additionally, we observed similar guide dependent effects on editing (Fig 2D left, right) as seen in figure 1, further confirming these results are dependent on direct genome modification, and not some other non-nuclease mediated effects or background from our assay. Finally, we quantified average editing rates for the 5’ and 3’ regions and INDEL rates surrounding the cutsites, as shown in Fig 2E and F. Again, INDEL rate largely shows good correlation with editing rate, with Cas9-HR consistently showing reduced INDELs compared to Cas9.

### Two guide editing strategies can perform long-range editing over ∼8kb genomic region

At that point, we were very confident in our ability to perform edits over ∼4kb regions, and wanted to test what the limits were for our long-range editing strategy by attempting to perform edits over a larger window. We chose to focus on an almost ∼8kb region including the ∼7.5kb exon 30 of the ApoB gene, which contains roughly ∼50% of all likely pathogenic and confirmed pathogenic mutations in the ApoB gene. To test this, we created a repair template ∼9kb in length, consisting of 17 individual mutations spanning ∼8kb (Fig 3A), and again tested four different two-guide combinations. Unsurprisingly, given the size of repair template, editing rates were the lowest seen so far, ranging from 0.2-0.8% depending on guide and editing position (Fig 3B, left). Unlike for Figures 1 or 2, Cas9-HR was not able to drive significant editing above background levels across the whole editing window (Fig 3B, right), demonstrating that different nucleases might show different efficiencies depending on genomic region targeted. Again, we saw guide dependent effects on editing rates, though this time with more pronounced spikes around the cutsites compared to the other two loci we tested. Finally, editing and INDEL rates were quantified for all guides as shown previously (Fig 3C,D). Although editing was not optimal, Cas9-HR still had reduced INDELs compared to Cas9, and all guide combinations showed good INDEL production. These results demonstrate that the editing window can be extended to over 8,000 base-pairs (and likely longer), as endogenous homologous recombination and linkage takes place over 100,000s of base-pairs.

**Figure 3:**
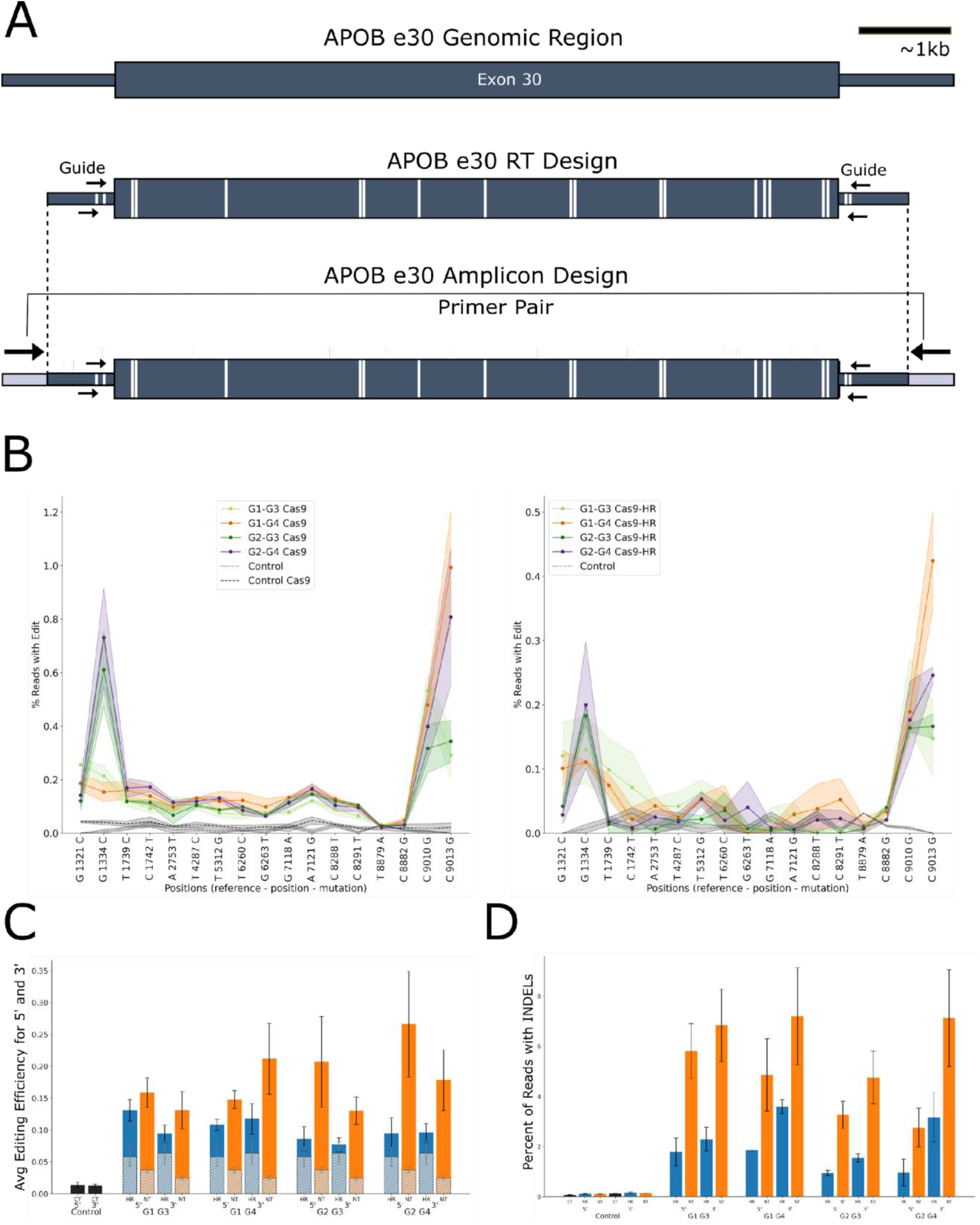
Two Guide strategies can drive long-range editing of ∼9kb region of ApoB exon 30. **3A**: Diagram showing the structure of the ApoB exons 30 genomic region. The regions denoting the repair template are shown by dark blue shading, vertical white bars denote specific mutations, short black arrows (guides used and orientation), bottom shows integrated repair template with primers used for amplification from genomic DNA (large black arrows and grey shaded region respectively). **3B**: Graph showing editing rates across ∼9kb of genomic DNA using various guide pairs. Left, Cas9; right, Cas9-HR. **3C**: Average editing of various double primer pairs divided into 5’ and 3’ regions. Blue, Cas9-HR; Orange, Cas9; Black, untransfected control; Grey hatched bars, Cas9-HR or Cas9 and no guide RT controls. **3D**: Graph showing the average percentage of reads containing INDELs around +/- 25 of 5’ and 3’ cut sites for single and double guide combinations. Error bars show SEM, n=2-3.

### Long-range editing strategies are compatible with other nucleases

Next, we wanted to test whether our strategies were compatible with nucleases other than the most commonly used Streptococcus pyogenes Cas9 (SpCas9). Therefore, we decided to use Staphylococcus Lugdunensis Cas9 (SluCas9), which has a very similar PAM (NNGG compared to NGG) to SpCas9, allowing us to use virtually the same sequences as our previously identified guides. Though SluCas9’s PAM sequence are shifted by one base-pair, SluCas9 is significantly smaller (∼3kb, 1,053 amino acids vs ∼4kb, 1,368 amino acids for SpCas9) and would enable AAV based editing^20,21^. We also switched from a two piece PCR strategy used in Figure 1 (Figure 4A), to a single PCR strategy similar to those used in Figures 2 and 3. Finally, we also created a SluCas9-HR fusion protein which we then tested along with SluCas9 for the ability to edit Factor 8 exon 14 and PAH exons 6-8, using 1bp shifted versions of the optimal guide pairs (G2+G5 and G2+G4 respectively) identified previously.

**Figure 4:**
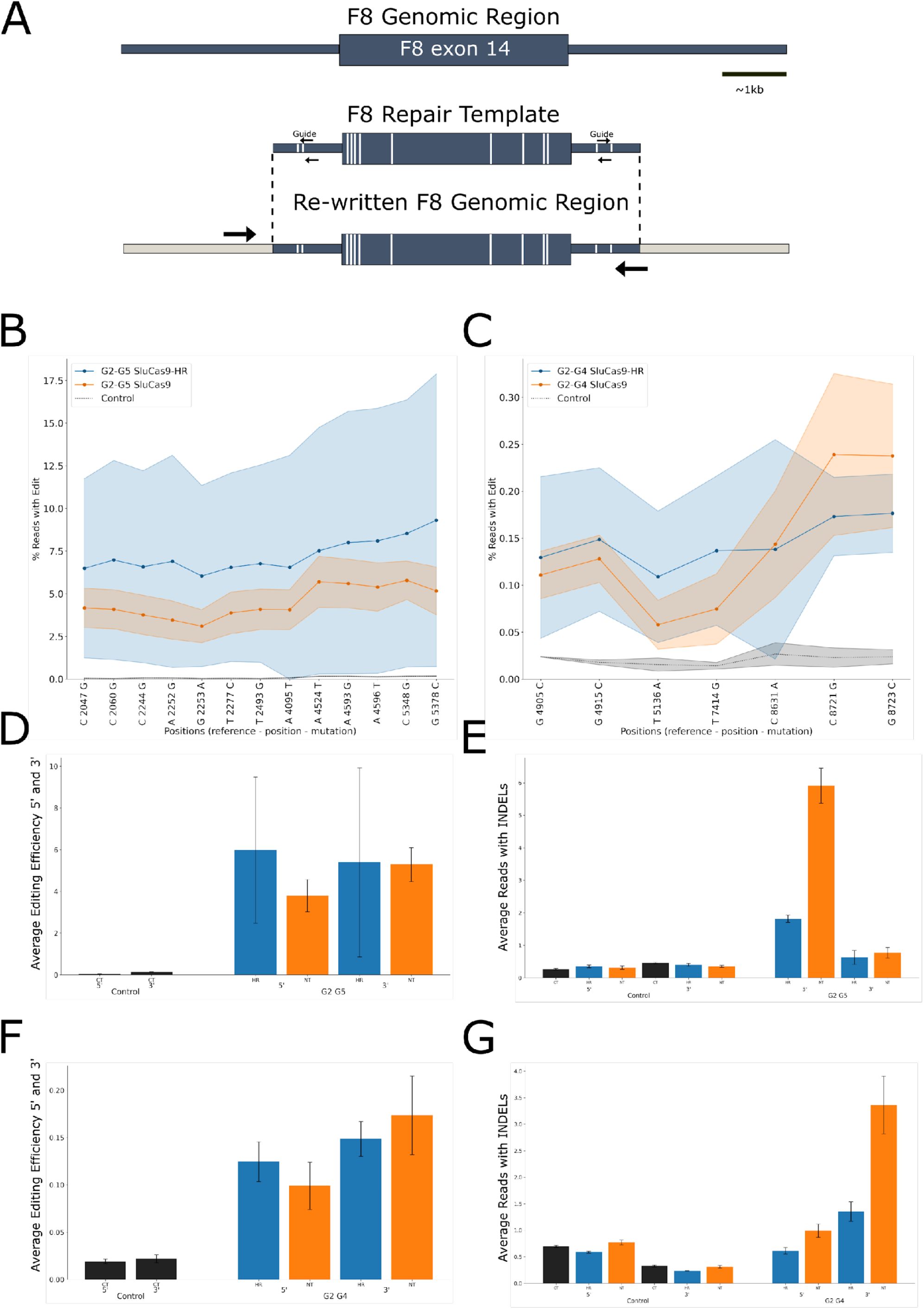
SluCas9, a SpCas9 orthologue, and SluCas9-HR also enable long-range editing. **4A**: Diagram showing the structure of the ApoB exons 30 genomic region. The regions denoting the repair template are shown by dark blue shading, vertical white bars denote specific mutations, short black arrows (guides used and orientation), bottom shows integrated repair template with primers used for amplification from genomic DNA (large black arrows and grey shaded region respectively). **4B**: Graph showing background subtracted editing rates across ∼3.3kb of genomic DNA using G2+G5 F8 guide pair. Blue, SluCas9-HR; Orange, SluCas9; Black, untransfected control. **4C**: Graph showing background subtracted editing rates across ∼4kb of genomic DNA using G2+G4 PAH guide pair. Blue, SluCas9-HR; Orange, SluCas9; Black, untransfected control. **4D**: Average editing of various double primer pairs divided into 5’ and 3’ regions. Blue, Cas9-HR; Orange, Cas9; Black, untransfected control; Grey hatched bars, Cas9-HR or Cas9 and no guide RT controls. **4E**: Graph showing the average percentage of reads containing INDELs around +/- 25 of 5’ and 3’ cut sites for single and double guide combinations. Error bars show SEM, n=3. **4F**: Average editing of various double primer pairs divided into 5’ and 3’ regions. Blue, Cas9-HR; Orange, Cas9; Black, untransfected control; Grey hatched bars, Cas9-HR or Cas9 and no guide RT controls. **4G**: Graph showing the average percentage of reads containing INDELs around +/- 25 of 5’ and 3’ cut sites for single and double guide combinations. Error bars show SEM, n=3.

Background subtracted results are shown in Figure 4B and 4C respectively, with both nucleases able to perform long-range editing across the previously established windows. Average editing efficiencies across F8 exon 14 (Fig 4D) and PAH exons 6-8 (Fig 4F) again correlated well with INDEL levels (Fig 4E and 4G, respectively). As with SpCas9-HR, SluCas9-HR showed significantly reduced levels of INDELs compared to SluCas9, though guide efficiency (and INDEL patterns produced, as previously reported; data not shown) differ between the two species of Cas9. These results confirm that chunk editing can be performed by multiple different nucleases, and likely by virtually any targetable nuclease.

### Non-DSB based nickase strategies also enable long-range editing

Finally, we wanted to test whether chunk editing can be used for non-DSB based applications. We designed two different approaches, both using our previously established F8 chunk editing assays. First, we created four different nickase variants: Cas9 D10A, Cas9-HR D10A, Cas9 H840A, Cas9-HR H840A. Nickase mutants D10A and H840A cut opposite strands of DNA^22,23^, and we hypothesized resection of the individual nicks via the exonuclease domain of Cas9-HR could increase rates of chunk editing. Finally, since we were still using plasmid-based transfection and thus were unable to simultaneously use D10A and H840A nucleases, we decided to take advantage of alternate orientation of previously identified guide pairs G2-G5 (same strand), and G2-G6 (staggered strands), and by using either D10A or H840A variants we could test all various permutations of strand nicking (Fig 5A). We first tested same-strand targeting of both guides (G2-G5) with results and editing efficiencies predictably low, with most not reaching above background levels (Figure 5B, quantified in 5D). Differences between D10A and H840A were seen, with H840A showing slightly higher editing rates than D10A. Cas9-HR did not seem to have any appreciable effect, instead, we saw staggered or same-stand strand nickase targeting having much larger effects. Next, we tested whether staggard strand approaches would increase long-range editing efficiency. As seen in Figure 5C and quantified in 5E, staggered targeting dramatically increases long-range editing efficiency with rates reaching >10% for certain positions. As with same strand targeting, the identity of the nickase (D10A or H840A) had a much stronger effect on editing efficiency than whether Cas9 or Cas9-HR were used. Finally, we noted via analysis of INDELs around the cutsites that all versions of nickases did not create INDELs over background. Interestingly, the fact that both staggered orientations were able to perform long-range editing and that long-range nicks seemingly don’t generate INDELs over background levels point to the fact that long-range nickase based editing functions differently than short range nickase based editing approaches. Finally, these results demonstrate that non-DSB based long-range editing is possible, and combined with previous experiments demonstrate the incredible flexibility of our novel long-range editing approach.

**Figure 5:**
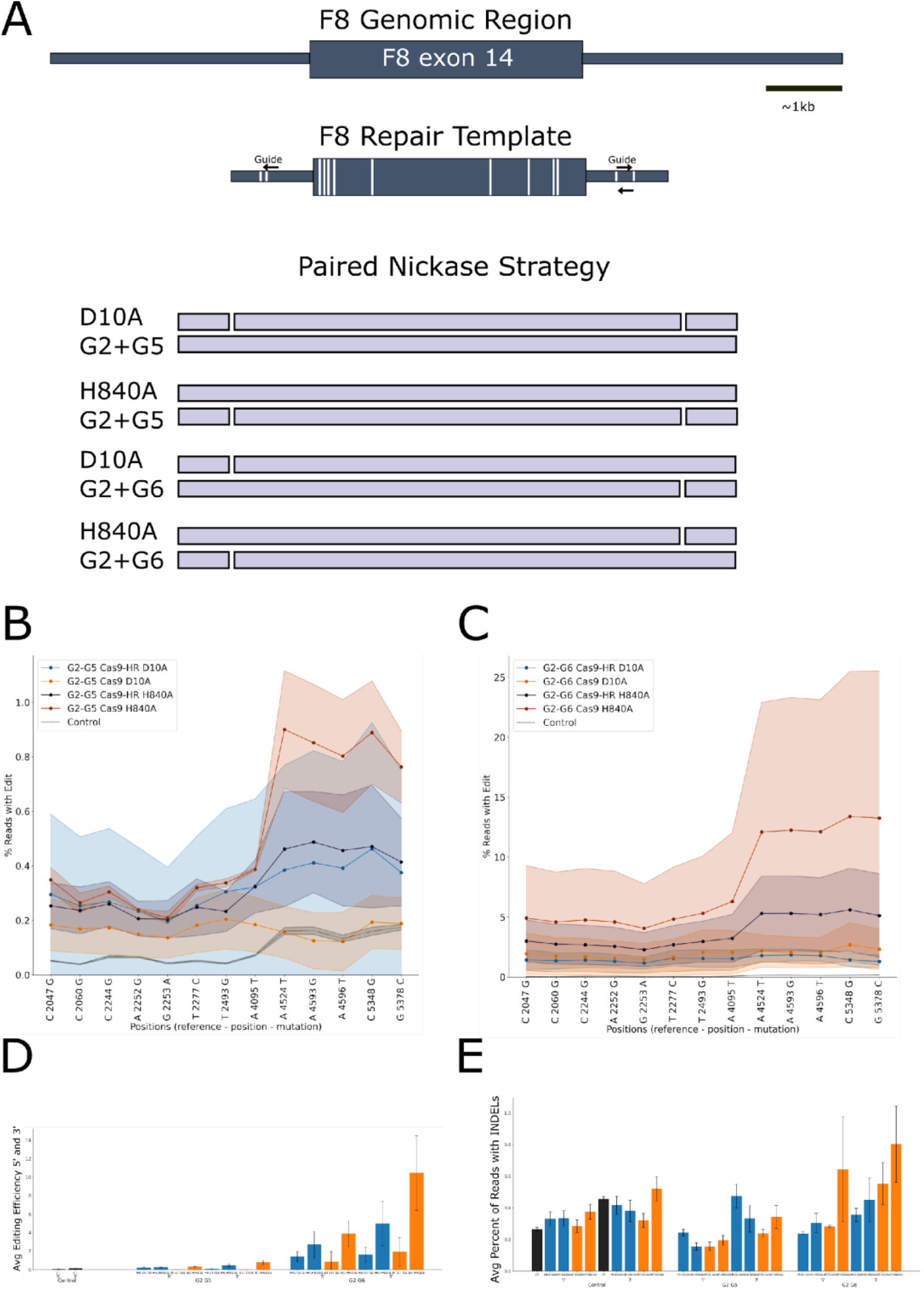
Long-range editing is compatible with nickase based non-DSB approaches. **5A**: Diagram showing diagram of various nickase and guide combinations to allow testing of all 4 types of editing permutations. **5B**: Graph showing background subtracted editing rates across ∼3.3kb of genomic DNA using G2+G5 F8 guide pair. Blue, Cas9-HR D10A; Orange, Cas9 D10A; Dark Blue, Cas9-HR H840A; Maroon Cas9 H840A; Black, untransfected control. **5C**: Graph showing background subtracted editing rates across ∼3.3kb of genomic DNA using G2+G6 F8 guide pair. Blue, Cas9-HR D10A; Orange, Cas9 D10A; Dark Blue, Cas9-HR H840A; Maroon Cas9 H840A; Black, untransfected control. **5D**: Average editing of various double primer pairs divided into 5’ and 3’ regions. Blue, Cas9-HR; Orange, Cas9; Black, untransfected control; Grey hatched bars, Cas9-HR or Cas9 and no guide RT controls. **5E**: Graph showing the average percentage of reads containing INDELs around +/- 25 of 5’ and 3’ cut sites for single and double guide combinations. Error bars show SEM, n=2-3.

## Discussion

These experiments have demonstrated that it is possible to use dual guides to make discrete and simultaneous rewrites over 1000s of base-pairs. We first demonstrated that either Cas9-HR or Cas9 can perform long rewrites in H1299 cells, and that these edits roughly correlated to guide cutsite INDEL rates. Next, we demonstrated that chunk editing was functional at an additional locus, and further explored how different guides affected chunk editing efficiency across the entire editing window. We also showed that chunk editing is functional in HEK293 cells, and that the editing pattern across the window was remarkably similar to that of H1299 cells. Further experiments at an additional locus, ApoB, showed that the editing window can be extended to a remarkable ∼7.5kb, though at lower efficiency compared to Factor 8 or PAH. We also demonstrated that multiple nucleases can perform long range rewrites, as both SluCas9 and SluCas9-HR were functional at both Factor 8 and PAH. Finally, we demonstrated dual nickase based strategies can be used to achieve remarkably high rates of long-range editing, with the best combinations yielding efficiencies of 5-10% across the whole window.

Initially, we were surprised to see that long range rewriting was possible with Cas9, as we expected Cas9-HR to uniquely facilitate this due to its enhanced HDR vs NHEJ properties. As expected, we did see a consistent and significant reduction in INDEL rates at virtually every site and guide tested, though a few were more modest. Additionally, we saw that Cas9-HR vs Cas9 editing efficiency significantly varied site-by-site, indicating that the use of Cas9-HR vs Cas9 is an additional variable to optimize. It will also be interesting to assay additional sites and cell types in the future.

Most surprisingly, opposite strand nickase-based chunk editing seems very effective, with efficiencies of 5-10% for the optimal configuration. We tried two different types of nickase strategies: same strand and opposite strand. Unsurprisingly, same-strand editing was largely ineffective with most nickases showing minimal to no editing across the window, though H840A nickases did show modest editing. However, opposite strand strategies showed remarkably improved rates, especially for H840A. We were initially surprised that nicks over such a large distance were able to be used for our chunk editing experiments, however, single nick based insertion of hundreds of base-pairs has been observed by both our in-house experiments (data not shown) as well as by others^25–27^. It will be very interesting to explore further how nickase based chunk editing operates in the future.

Practically, we think further development of long-range editing could have significant clinical applications, particularly in the rare disease space. Currently, the vast majority of corrective approaches are limited to single therapeutic correcting a single mutation (Fig 6A). While this is effective in specific situations, the vast majority of human monogenic rare diseases have multiple documented pathogenic variants (Fig 6B), and this dramatically limits the practical use of these strategies to address rare diseases with diverse mutational profiles (Fig 6C). We believe long-range editing could dramatically change this paradigm by enabling an approach where a single therapeutic could treat many different mutations. A long-range editing approach as described above could significantly increase both the scientific and financial viability of corrective approaches given the significant time, effort and money needed to bring a therapeutic to market.

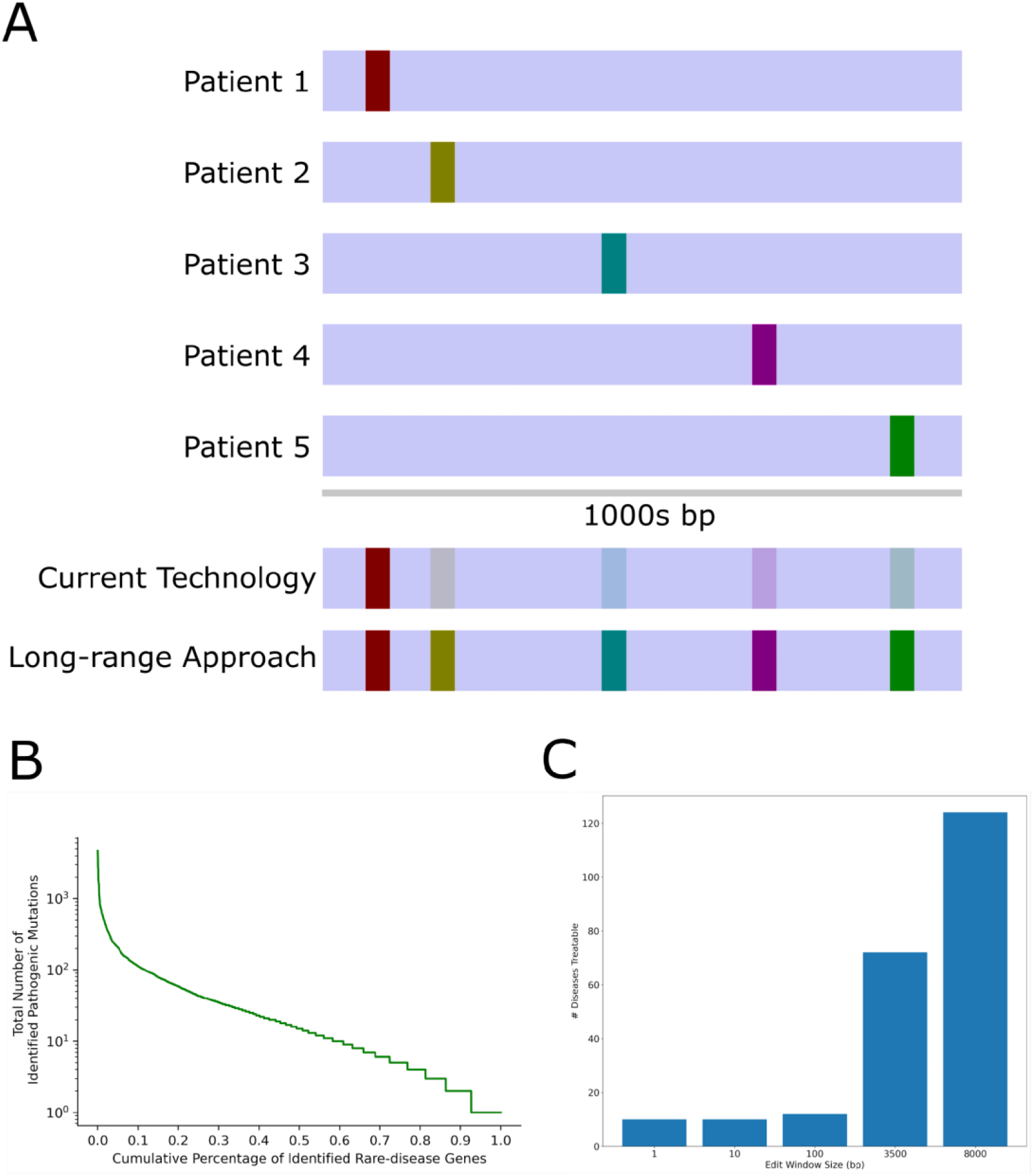
Implications of long-range rewriting on rare disease therapeutic approaches. **6A**: Top diagram shows an example monogenic rare disease caused by different mutations in different patients. Bottom diagram shows current limitation of corrective based editing approaches (generally limited to very short windows and single mutations), where a single therapeutic can treat a single mutation. Long-range editing, however, has the potential to treat many different mutations with a single therapeutic. Colored vertical bars denote various pathogenic mutations, blue shaded bar is the example genomic region, faded bars denote untreated mutations. **6B**: Majority of human monogenic rare diseases are caused by diverse mutations. Diagram showing the total number of pathogenic mutations identified for >3,000 genes with known links to rare diseases. Cumulative percentage of genes is shown on the x-axis, with the y-axis (log scale) showing the total number of known pathogenic mutations for each gene. Data adapted from ClinVar and Orphanet. **6C**: Long-range editing and increased editing window size dramatically increases the number of diseases with >300 patients which can be treated with a single therapeutic. Bar graph showing the effect of increasing the window size of on the number of indications with estimated >300 patients. Y-axis denotes total indications with estimated patient populations >300 for the given editing window size, as denoted by the x-axis. Data adapted from ClinVar.

While these experiments have demonstrated long range rewrites can be performed using a variety of strategies, editing rates are still relatively low as little optimization has been done to date. In particular, our experiments generally generated low INDEL levels, likely due to use of plasmids. Experimenting with different DNA repair template types, guide and nuclease delivery modalities will likely dramatically increase editing rates, as has been seen for insertional based applications^24^. Additionally, it is apparent from these experiments that the rules determining chunk editing efficiency are more complex than single guide-based applications. For example, PAH G3 is ineffective in both INDEL and editing efficiency when used alone, however pairing it with a stronger guide such as PAH G2 results in editing, though at reduced efficiency. Future experiments should help to illuminate how INDEL production, editing window size, and local chromatin state amongst other factors all affect long range rewriting efficiency. In conclusion, we have demonstrated that long range rewriting is a powerful new technique which has the potential to dramatically impact a variety of therapeutic and basic R&D applications.

---

**Figure 1S:**
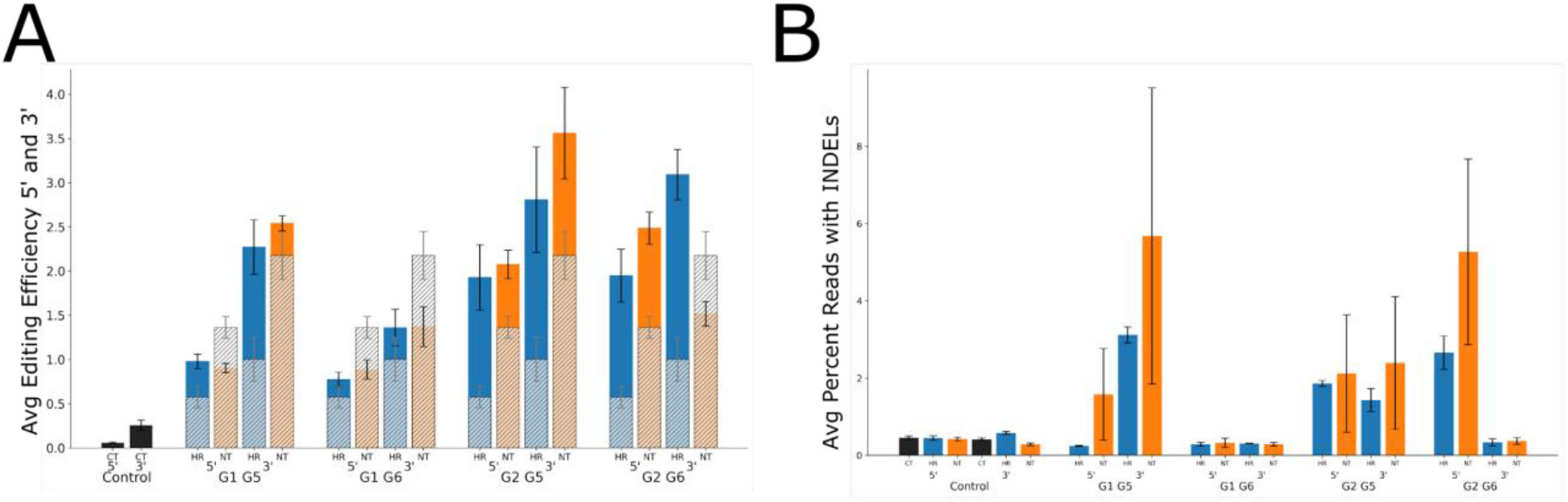
Long-range editing efficiency generally correlates with guide INDEL generation. At least one effective guide is necessary to enable chunk editing. Fig S1A: Graph showing the average editing efficiency across 5’ (positions 2047-2493) and 3’ (4095-5378) of Factor 8 exon 14. Solid bars represent denote guide and nuclease pairs plus repair template, with semi-transparent shaded bars denoting controls using corresponding nuclease and repair template but lacking guides. Black bars to left show background levels of editing in untransfected cells. n=2-3 per experiment, error bars represent SEM. Fig S1B: Bar graph showing average percent of reads containing INDELs for each of the guide (below), nuclease (Cas9-HR blue, Cas9 orange) and 5’ and 3’ ends as denoted by the x-axis legend, replicates and error bars are the same as in FigS1A.

**Figure 2S:**
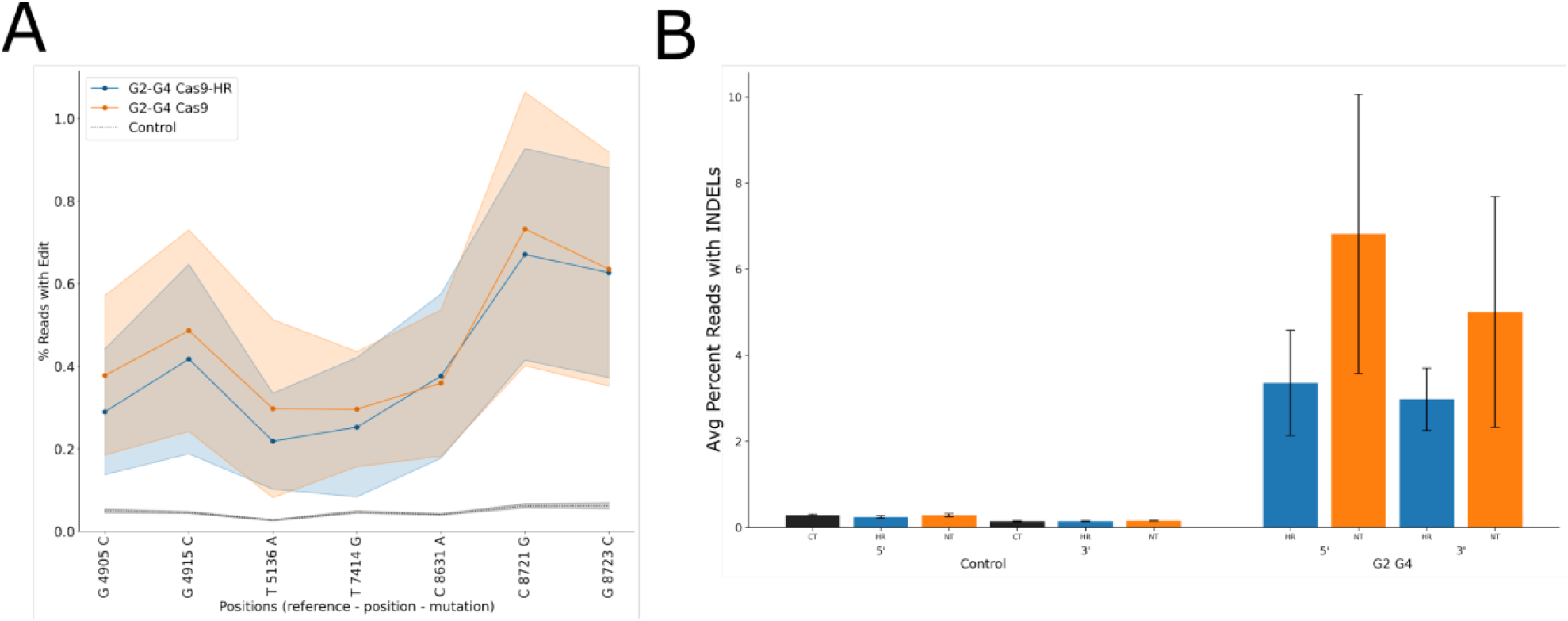
Long-range editing in HEK 293 cells shows similar editing and INDEL patterns compared to H1299 cells. Fig S2A: Graph showing chunk editing in HEK293 cells showing background subtracted (Cas9-HR+RT and Cas9+RT respectively) editing rates for both Cas9-HR (blue) and Cas9 (orange) compared to non-transfected cells, n=3, error bars=SEM. Fig S2B: Graph showing the Average percentage of reads containing INDELs around +/- 25 of 5’ and 3’ cut sites for G2-G4 guide combinations, n=3, error bars=SEM.

